# Growth plate resting zone chondrocytes acquire transient clonal competency upon Hedgehog activation and efficiently transform into trabecular bone osteoblasts

**DOI:** 10.1101/2023.05.31.543069

**Authors:** Shion Orikasa, Yuki Matsushita, Michael Fogge, Koji Mizuhashi, Naoko Sakagami, Wanida Ono, Noriaki Ono

**Author notes:** **Correspondence:** Noriaki Ono.

## Abstract

The resting zone of the postnatal growth plate is organized by slow-cycling chondrocytes expressing parathyroid hormone-related protein (PTHrP), which include a subgroup of skeletal stem cells that contribute to the formation of columnar chondrocytes. The PTHrP–indian hedgehog (Ihh) feedback regulation is essential for sustaining growth plate activities; however, molecular mechanisms regulating cell fates of PTHrP^+^ resting chondrocytes and their eventual transformation into osteoblasts remain largely undefined. Here, in a mouse model, we utilized a tamoxifen-inducible *PTHrP-creER* line with *Patched-1* (*Ptch1*) floxed and tdTomato reporter alleles to specifically activate Hedgehog signaling in PTHrP^+^ resting chondrocytes and trace the fate of their descendants. Hedgehog-activated PTHrP^+^ chondrocytes formed large concentric clonally expanded cell populations within the resting zone (‘*patched roses*’) and generated significantly wider columns of chondrocytes, resulting in hyperplasia of the growth plate. Interestingly, Hedgehog-activated PTHrP^+^ cell-descendants migrated away from the growth plate and eventually transformed into trabecular osteoblasts in the diaphyseal marrow space in the long term. Therefore, Hedgehog activation drives resting zone chondrocytes into transit-amplifying states as proliferating chondrocytes and eventually converts these cells into osteoblasts, unraveling a novel Hedgehog-mediated mechanism that facilitates osteogenic cell fates of PTHrP^+^ skeletal stem cells.

## Introduction

The growth plate – a thin disk-like cartilage located at the edges of long bones – provides the principal driving force for postnatal bone growth (1). Structurally, the growth plate is composed of three morphologically distinct layers of resting, proliferating and hypertrophic zones with characteristic columns of chondrocytes with clonal origins (2, 3). The growth plate is an essential structure for endochondral bone formation, the process by which the transient cartilage is gradually replaced by the bone (1, 2). The resting zone located at the top of the postnatal growth plate houses slow-cycling chondrocytes expressing parathyroid hormone-related protein (PTHrP) (4), which provide a source of all other chondrocytes within the growth plate. These “*resting*” chondrocytes enter the cell cycle through asymmetric divisions, become proliferating chondrocytes, differentiate into post-mitotic prehypertrophic chondrocytes that express Indian hedgehog (Ihh), become hypertrophic chondrocytes at the bottom of the growth plate, and eventually undergo either cell death due to apoptosis or transformation into trabecular bone osteoblasts in the primary spongiosa.

The resting zone of the postnatal growth plate contains populations of stem-like cells that give rise to clones of proliferating chondrocytes and determine their orientation, as first identified by autologous surgical transplantation experiments in rabbits (5) and later confirmed by in vivo genetic lineage-tracing experiments (6). The resting zone is organized by the stem cell subgroups that play important roles in long-term tissue renewal, including PTHrP^+^ (6), FoxA2^+^ (7) cells and other slow-cycling cells that are maintained in a Wnt-inhibitory environment (8), collectively referred to as epiphyseal skeletal stem cells (9)(10). Signals from the epiphyseal stem cell niche – the secondary ossification center (SOC), composed of a variety of bone cells such as osteoblasts, osteocytes, hypertrophic chondrocytes, and hematopoietic cells – play important roles in forming these stem cells within the resting zone (11).

The PTHrP–Ihh negative feedback system is essential for maintaining activities of the growth plate. PTHrP functions as a forward signal from undifferentiated chondrocytes in the resting zone, stimulating proliferation of chondrocytes in the adjacent proliferating zone through binding to its cognate receptor PTH1R and inhibiting their terminal differentiation into hypertrophic chondrocytes (12–14), whereas Ihh functions as a reverse signal from terminally differentiated chondrocytes in the prehypertrophic zone, stimulating proliferation of chondrocytes in the adjacent proliferating zone and their differentiation into columnar chondrocytes, while increasing PTHrP expression in the resting zone (15–17). The functional interaction between PTHrP and Ihh likely plays important roles in regulating skeletal stem cell behaviors. In Hedgehog signaling, Ihh binds to its cognate receptor Patched 1 (PTCH1) and relieves its inhibitory function on the G-protein coupled receptor Smoothened (SMO), leading to activation of Gli transcription factors (18). Ihh released from prehypertrophic chondrocytes within the cartilage template is essential for the formation of osteoblast precursors in the perichondrium (19–23). Gli transcription factors promote osteoblast differentiation by activating BMP functions and inducing early osteoblast differentiation in a Runx2-independent manner (24, 25). Therefore, Hedgehog signaling plays important roles in osteoblast differentiation.

A subset of PTHrP^+^ resting chondrocytes represents a unique type of skeletal stem cells which are initially unipotent and later acquire multipotency at the post-mitotic stage. Hypertrophic chondrocytes transform into osteoblasts and CXCL12-abundant reticular (CAR) cells as these cells exit from the growth plate (6). Chondrocyte-to-osteoblast transformation has been demonstrated by several key in vivo lineage-tracing studies (26–28). Importantly, our previous lineage-tracing study implies that the hypertrophic chondrocyte-to-osteoblast transformation is not an efficient process, as the descendants of PTHrP^+^ resting chondrocytes contribute to only a small number of osteoblasts and CAR cells in the adult bone marrow (6). Molecular mechanisms regulating this dynamic transformation remain largely unknown.

In this study, we hypothesized that Hedgehog signaling facilitates osteogenic cell fates of PTHrP^+^ resting chondrocytes through multiple mechanisms. We addressed this question by an in vivo functional cell lineage analysis using a *PTHrP-creER* line that we reported in our previous study (6). This line can exclusively mark PTHrP^+^ chondrocytes in the resting zone of the postnatal growth plate upon tamoxifen injection, without marking any other cell types in long bones. Capitalizing on this highly cell type-specific genetic tool, our goal was to define the long-term cell fate of PTHrP^+^ resting chondrocytes when Hedgehog signaling was specifically manipulated in these cells using *Ptch1*-floxed or *Smo*-floxed alleles, to activate or deactivate Hedgehog signaling, respectively. Our *PTHrP-creER*-based approach sets it apart from preceding studies utilizing other inducible *creER* lines such as *Prrx1-creER* (29) or *Col2a1-creER* (17, 30), both of which mark a wide variety of cells in the osteoblast and chondrocyte lineages.

Our findings overall demonstrate that Hedgehog-activated PTHrP^+^ resting chondrocytes transiently acquire clonal competency within the resting zone and induce hyperplasia of the growth plate, and subsequently, their descendants migrate away from the growth plate and efficiently transform into trabecular bone osteoblasts in the diaphyseal marrow space. Our study strengthens the link from PTHrP^+^ chondrocytes in the resting zone to osteoblasts in the trabecular bone based on a functional genetic approach, unraveling a novel Hedgehog-mediated mechanism to regulate ultimate osteogenic cell fates of PTHrP^+^ skeletal stem cells.

## Results

### 1. PTHrP^+^ resting chondrocytes lose quiescence and establish “patched roses” upon Hedgehog activation

We employed a *PTHrP-creER* line that can specifically mark a group of PTHrP^+^ chondrocytes in the resting zone upon tamoxifen injection (6); importantly, this line does not mark any other growth plate chondrocytes in different layers, osteoblasts/cytes or bone marrow stromal cells, enabling a highly cell type-specific lineage analysis of PTHrP^+^ resting chondrocytes in the context of physiological endochondral bone growth occurring during postnatal stages (Fig.1A,B). Using *PTHrP-creER*, we conditionally deleted the Hedgehog cognate receptor PTCH1 using *Ptch1* floxed alleles (31), deletion of which causes ligand-independent activation of Smoothened (SMO) signaling, an essential downstream effector of Hedgehog ligands (18, 32, 33). We performed in vivo functional cell-lineage analysis of Hedgehog-activated descendants of PTHrP^+^ resting chondrocytes in a mosaic environment, by introducing a *cre*-responsive *R26R-tdTomato* reporter (Fig.1C). *PTHrP-creER; Ptch1^fl/+^; R26R^tdTomato^*(PTHrP-Ptch Control, PTHrP^CE^-P6 cells) and *PTHrP-creER; Ptch1^fl/fl^; R26R^tdTomato^* (PTHrP-Ptch cKO, PTHrP^CE^ΔPtch-P6 cells) mice were pulsed at P6 and analyzed at P14 and P21 to trace the fate of PTHrP^CE^-P6 and PTHrP^CE^ΔPtch-P6 cells by tdTomato epifluorescence.

**Figure 1.**
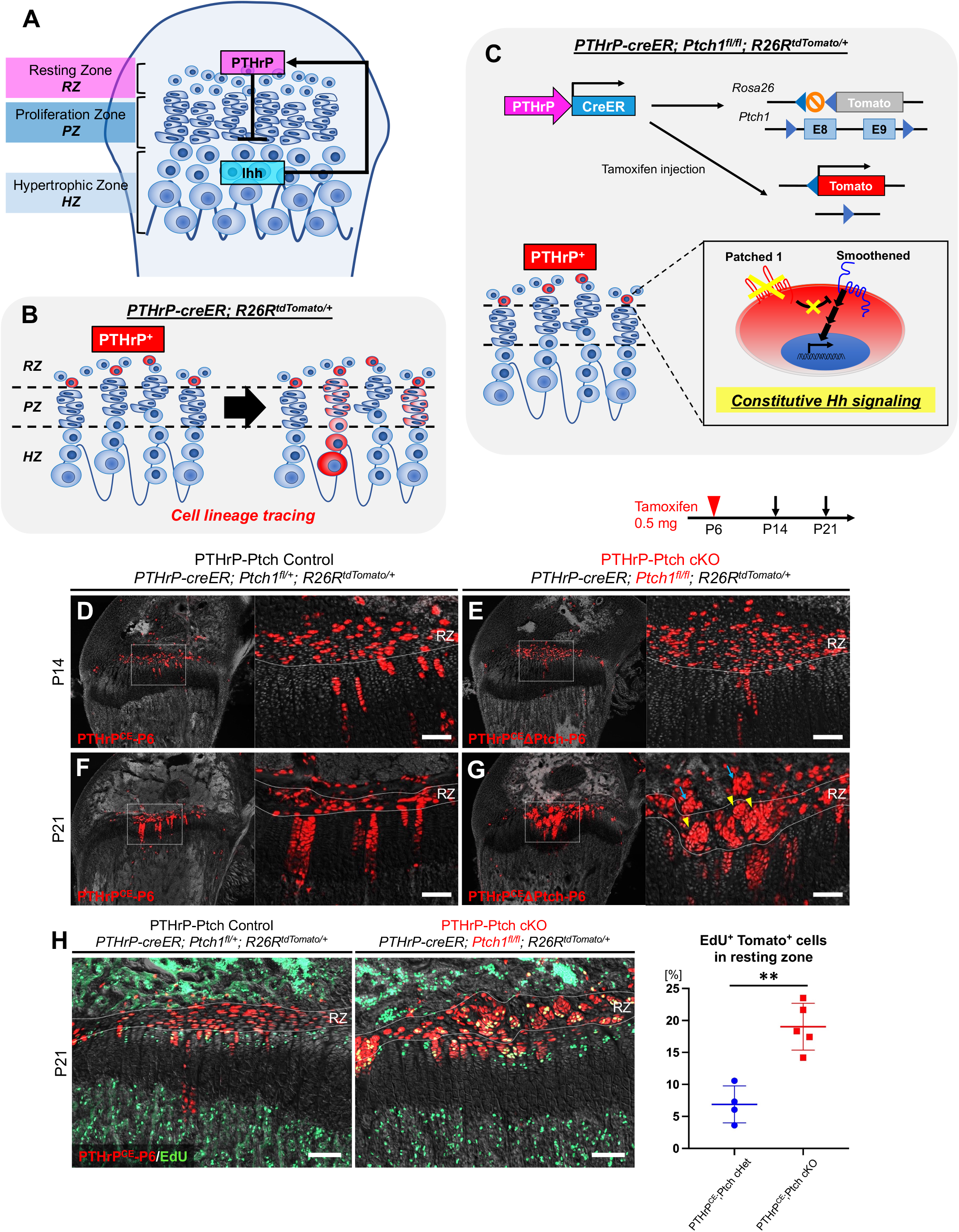
PTHrP^+^ resting chondrocytes lose quiescence and establish “patched roses” upon Hedgehog activation. **(A)** Postnatal growth plate structure with PTHrP–Ihh negative feedback system. **(B)** In vivo lineage-tracing of PTHrP^+^ resting chondrocytes using *PTHrP-creER; R26R^tdTomato/+^* bigenic mice. lineage-tracing approach. **(C)** Constitutive Hedgehog activation in PTHrP^+^ resting chondrocytes using *PTHrP-creER; Ptch1^fl/fl^; R26R^tdTomato^* trigenic mice. **(D-G)** Histological images of growth plates, *PTHrP-creER; Ptch1^fl/+^; R26R^tdTomato^* (PTHrP-Ptch Control, D,F) and *PTHrP-creER; Ptch1^fl/fl^; R26R^tdTomato^* (PTHrP-Ptch cKO, E,G) distal femur at P14 (D,E) and P21 (F,G) (pulsed at P6). RZ: resting zone, red: tdTomato, gray: DAPI. Scale bars: 100 μm. **(H)** Cell proliferation in *PTHrP-creER; Ptch1^fl/+^; R26R^tdTomato^*(Control) and *PTHrP-creER; Ptch1^fl/fl^; R26R^tdTomato^*(PTHrP-Ptch cKO) distal femur growth plate at P21 (pulsed at P6). EdU was administered 3 h before analysis. (Right panel): Quantification of EdU^+^tdTomato^+^ cells. *n*=4 (Control), *n*=5 (PTHrP-Ptch cKO) mice. Scale bars: 100 μm. \*\**p*<0.01, two-tailed, Mann-Whitney’s *U*-test. Data are presented as mean ± s.d.

PTHrP^+^ resting chondrocytes enter cell cycle and start to form columnar chondrocytes after ∼7 days of chase following a pulse at P6 (6). As expected, PTHrP^CE^-P6 cells formed short columns predominantly composed of less than 10 cells at P14 (Fig.1D). PTHrP^CE^ΔPtch-P6 cells behaved in a similar manner at this stage (Fig.1E), indicating that Hedgehog activation does not initially alter the behavior of PTHrP^+^ resting chondrocytes prior to the formation of the secondary ossification center. Upon further chase at P21, PTHrP^CE^-P6 cells formed long columns composed of more than 10 cells, which extended longitudinally and typically consisted of no more than two to three cells in width (Fig.1F). In contrast, PTHrP^CE^ΔPtch-P6 cells aberrantly expanded within the resting zone and formed “*patched roses*”, defined as concentric seemingly clonally expanded populations of PTHrP^+^ resting chondrocytes (Fig.1G). Evaluation of cell proliferation by EdU revealed that, while PTHrP^CE^-P6 cells predominantly remained quiescent in the resting zone, PTHrP^CE^ΔPtch-P6 cells became highly proliferative in the resting zone seemingly in an equal proportion to those in the proliferating zone (Fig.1H, left panel). Reflecting this observation, the percentage of EdU^+^ was significantly increased in Hedgehog-activated PTHrP^CE^ΔPtch-P6 cells compared to that of Control PTHrP^CE^-P6 cells (Fig.1H, right panel). Therefore, Hedgehog activation induces loss of quiescence and aberrant clonal expansion of PTHrP^+^ chondrocytes within the resting zone, leading to the formation of clonally expanded cell clusters or *patched roses*.

### 2. Hedgehog activation in PTHrP^+^ resting chondrocytes causes hyperplasia of the growth plate

PTHrP^+^ resting chondrocytes contribute to the entire length of columnar chondrocytes after one month of chase when pulsed at P6 (6). At P36, the growth plate of Control mice was entirely composed of vertically aligned columns of chondrocytes spanning over the three different layers across the growth plate (Fig.2A,B, left panels). In contrast, in PTHrP-Ptch cKO mice, the boundary between the resting and proliferating zones was obliterated, wherein deranged clusters of chondrocytes bulged outward from the center of the growth plate in large masses, resulting in hyperplasia and distortion of the growth plate (Figure 2A,B, right panels). Notably, unlike PTHrP^CE^-P6 Control cells, Hedgehog-activated PTHrP^CE^ΔPtch-P6 cells behaved in a significantly less linear manner, displaying deviation from the longitudinal axis associated with a loss of polarity (Fig.2C,D). Columns of PTHrP^CE^ΔPtch-P6 cells expanded laterally, resulting in a drastic increase in tdTomato^+^ columns within the PTHrP-Ptch cKO growth plate (Fig.2D).

**Figure 2.**
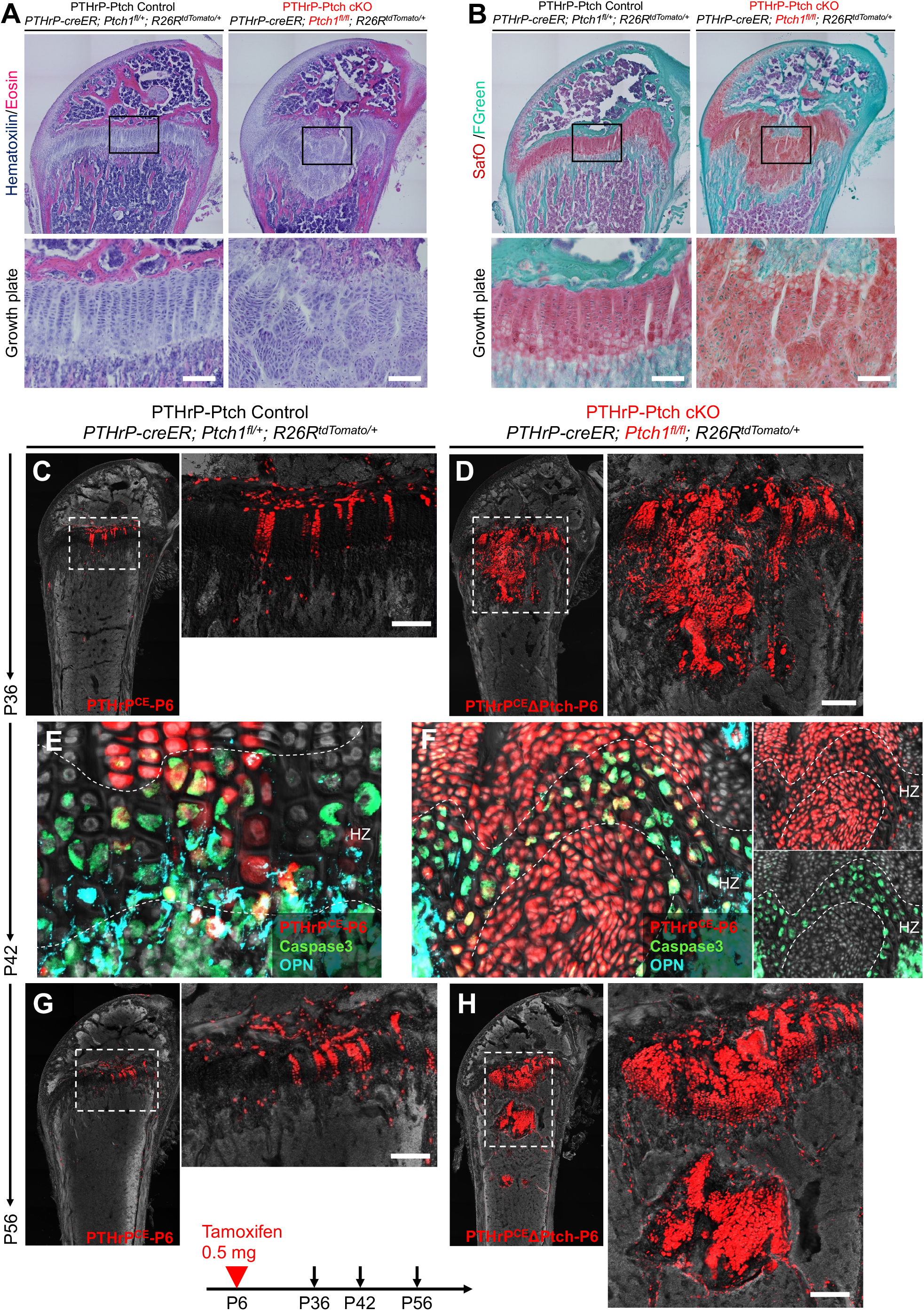
Hedgehog activation in PTHrP^+^ resting chondrocytes causes hyperplasia of the growth plate. **(A,B)** H&E (A) and Safranin O (B) staining of *PTHrP-creER; Ptch1^fl/+^; R26R^tdTomato^*(PTHrP-Ptch Control, left panels) and *PTHrP-creER; Ptch1^fl/fl^; R26R^tdTomato^* (PTHrP-Ptch cKO, right panels) distal femur at P36 (pulsed at P6). Scale bar: 100 μm. **(C,D)** Histological images of growth plates, *PTHrP-creER; Ptch1^fl/+^; R26R^tdTomato^* (PTHrP-Ptch Control, C) and *PTHrP-creER; Ptch1^fl/fl^; R26R^tdTomato^* (PTHrP-Ptch cKO, D) distal femur at P36 (pulsed at P6). Red: tdTomato, gray: DAPI. Scale bars: 100 μm. **(E, F)** Apoptosis in *PTHrP-creER; Ptch1^fl/+^; R26R^tdTomato^* (PTHrP-Ptch Control, E) and *PTHrP-creER; Ptch1^fl/fl^; R26R^tdTomato^* (PTHrP-Ptch cKO, F) hypertrophic zone at P42 (pulsed at P6). Immunostaining for Caspase 3 and osteopontin (OPN). HZ: hypertrophic zone. Green: Caspase 3 (apoptosis), red: tdTomato, light blue: OPN, Gray: DAPI.

We closely examined the hypertrophic zone at P42 to evaluate chondrocyte apoptosis. In PTHrP-Ptch Control mice, while a majority of PTHrP^CE^-P6 Control cells were positive for Caspase-3, we also noticed a small fraction of PTHrP^CE^-P6 cells that were negative for Caspase-3 in the hypertrophic layer, indicative of an escape from apoptosis (Fig.2E). In PTHrP-Ptch cKO mice, some Hedgehog-activated PTHrP^CE^ΔPtch-P6 cells were positive for Caspase-3 at the disorganized chondro-osseous junction (Fig.2F), while other PTHrP^CE^ΔPtch-P6 cells also appeared to be negative for Caspase-3, indicating that these cells can also escape from apoptosis.

We further examined the mutant growth plate at P56. In PTHrP-Ptch Control mice, PTHrP^CE^-P6 cells were primarily located within the growth plate, with a small number of these cells also observed in the bone marrow (Fig.2G). In contrast in PTHrP-Ptch cKO mice, clusters of PTHrP^CE^ΔPtch-P6 cells detached from the growth plate proper and formed a cartilaginous island in the metaphyseal marrow space (Fig.2H). Therefore, Hedgehog-activated PTHrP^+^ resting chondrocytes gain clonal competency and dominate the growth plate, resulting in hyperplasia of the growth plate, which detach from the growth plate and migrate toward the marrow space at later stages.

We next asked whether inactivation of Hedgehog signaling alters in vivo dynamics of PTHrP^+^ resting chondrocytes. To achieve this, we conditionally deleted *Smoothened* (*Smo*), a major downstream effector of Hedgehog signaling, using *Smo*-floxed alleles (34), and simultaneously traced the fates of these cells using an *R26R-*tdTomato reporter allele (Suppl.Fig.1A). Littermate triple transgenic mice with two corresponding genotypes – *PTHrP-creER; Smo^fl/+^; R26R^tdTomato^* (PTHrP-Smo Control, PTHrP^CE^-P6 cells) and *PTHrP-creER; Smo^fl/fl^; R26R^tdTomato^* (PTHrP-Smo cKO, PTHrP^CE^ΔSmo-P6 cells) mice were pulsed at P6 with tamoxifen and analyzed at P28. PTHrP^CE^ΔSmo-P6 cells established columnar chondrocytes in a seemingly normal pattern (Suppl. Fig.1A-C). However, the number of tdTomato^+^ columns in the PTHrP-Smo cKO was significantly less than that of the Control (Suppl. Fig.1D). The bone length and overall growth plate structure were unchanged in the PTHrP-Smo cKO (Suppl. Fig.1E and F). Therefore, inactivation of Hedgehog signaling in PTHrP^+^ resting chondrocytes slightly impairs their ability to establish columnar chondrocytes, without altering the overall morphology of the growth plate.

### 3. Hedgehog-activated PTHrP^+^ descendants transform into trabecular bone osteoblasts

Descendants of PTHrP^+^ resting chondrocytes contribute to the bone marrow and become osteoblasts and CXCL12-abandant reticular (CAR) stromal cells; this contribution increases for the first three months following a pulse at P6 (6). At P96, PTHrP^CE^-P6 Control cells were predominantly present in the growth plate as columnar chondrocytes with only a small number of cells present in the bone marrow (Fig.3A). In contrast, PTHrP^CE^ΔPtch-P6 cells contributed robustly to trabecular bone osteoblasts at the same stage. Strikingly, many PTHrP^CE^ΔPtch-P6 tdTomato^+^ cells were found to exist in the bone marrow and on the trabecular surface, contributing to an increase in the bone trabeculae within the PTHrP-Ptch cKO marrow space (Fig.3B,C). PTHrP^CE^ΔPtch-P6 cells located on the trabecular bone surface were closely associated with bone matrix proteins, osteopontin (OPN) and type I collagen (ColI). Interestingly, the PTHrP-Ptch cKO growth plate appeared largely normal without any hyperplasia, wherein PTHrP^CE^ΔPtch-P6 cells formed columnar chondrocytes in an organized pattern (Fig.3B). Therefore, Hedgehog-activated PTHrP^+^ descendants that migrate away from the growth plate efficiently transform into trabecular bone osteoblasts in adult stages.

**Figure 3.**
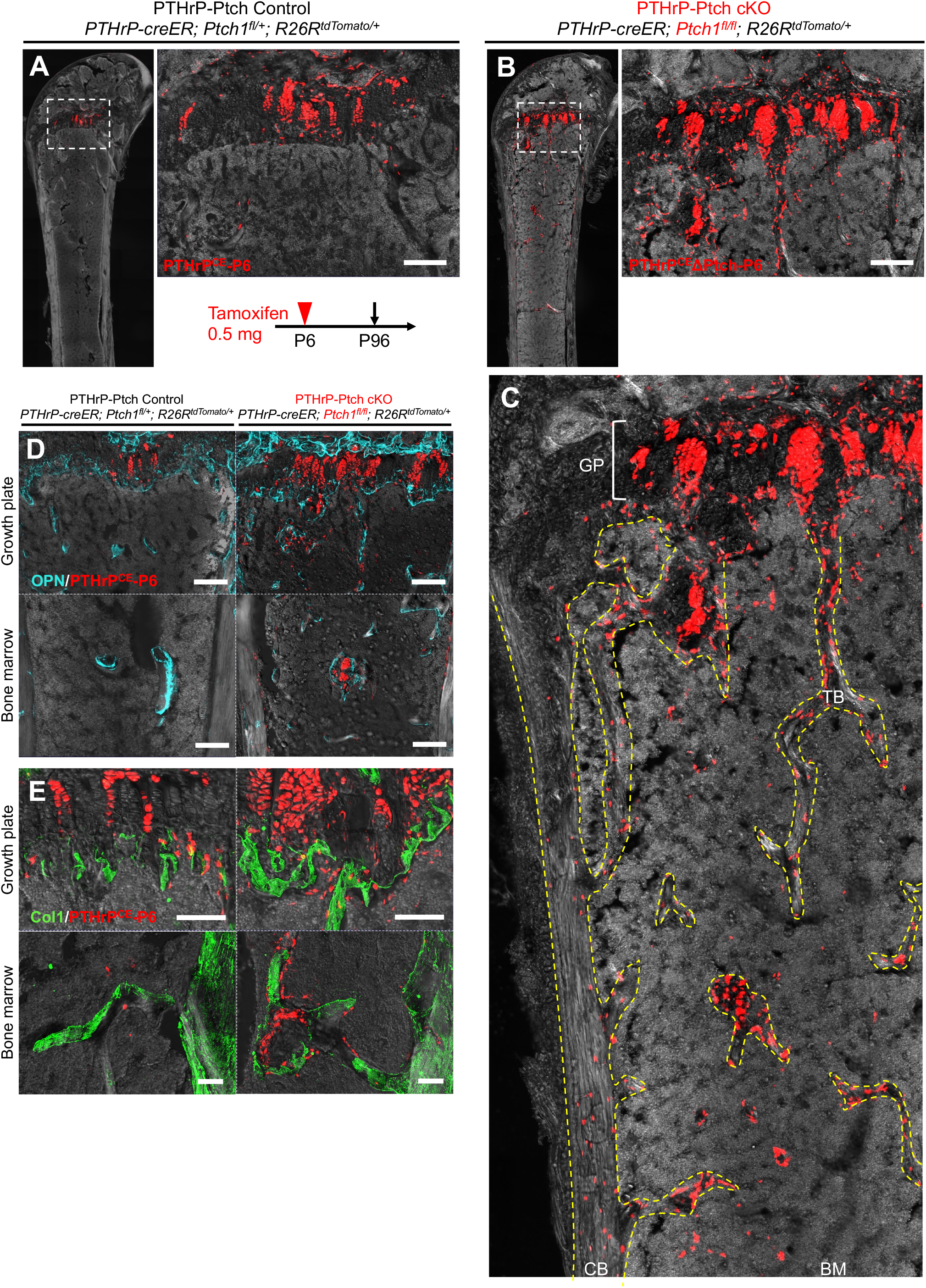
Hedgehog-activated PTHrP+ descendants transform into trabecular bone osteoblasts. **(A-C)** Histological images of distal femurs, *PTHrP-creER; Ptch1^fl/+^; R26R^tdTomato^* (PTHrP-Ptch Control, A) and *PTHrP-creER; Ptch1^fl/fl^; R26R^tdTomato^* (PTHrP-Ptch cKO, B,C) femurs at P96 (pulsed at P6). GP: growth plate, TB: trabecular bone, CB: cortical bone, BM: bone marrow. Yellow dotted lines: traces of bone surface. Red: tdTomato, gray: DAPI. Scale bars: 100 μm. **(D,E)** Bone matrix proteins. Immunostaining for OPN (D) and ColI (E). Scale bars: 200 μm (D), 100 μm (E).

### 4. Transient clonal competency of Hedgehog-activated PTHrP^+^ resting chondrocytes and their contribution to trabecular bone formation

To comprehensively define the kinetics of Hedgehog-activated PTHrP^+^ resting chondrocytes in the growth plate in an unbiased manner, we enumerated the number of lineage-traced PTHrP^CE^-P6 Control and PTHrP^CE^ΔPtch-P6 tdTomato^+^ cells across the time points ranging from P14 to P96 using Image J/Fiji-based cell quantification. First, we analyzed the number of tdTomato^+^ cells within the whole growth plate. In the PTHrP-Ptch Control growth plate, the number of PTHrP^CE^-P6 cells gradually increased up to P56, reaching a plateau thereafter (Fig.4A, blue bars). In contrast, in the PTHrP-Ptch cKO growth plate, PTHrP^CE^ΔPtch-P6 cells drastically and progressively increased between P21 and P36, then decreased at P56 and P70 in PTHrP-Ptch cKO mice (Fig.4B, red bars), reaching no statistical difference at P96. A similar pattern was observed when tdTomato^+^ cells were quantified in the resting zone: PTHrP^CE^-P6 cells gradually decreased within the resting zone up to P96 (6) (Fig.4B, blue bars), while PTHrP^CE^ΔPtch-P6 cells rapidly increased within the resting zone between P14 and P36, then decreased toward P96 (Fig.4B, red bars). These quantitative data corroborate the histological observation that PTHrP^CE^ΔPtch-P6 cells transiently gain clonal competency within the resting zone at P36.

**Figure 4.**
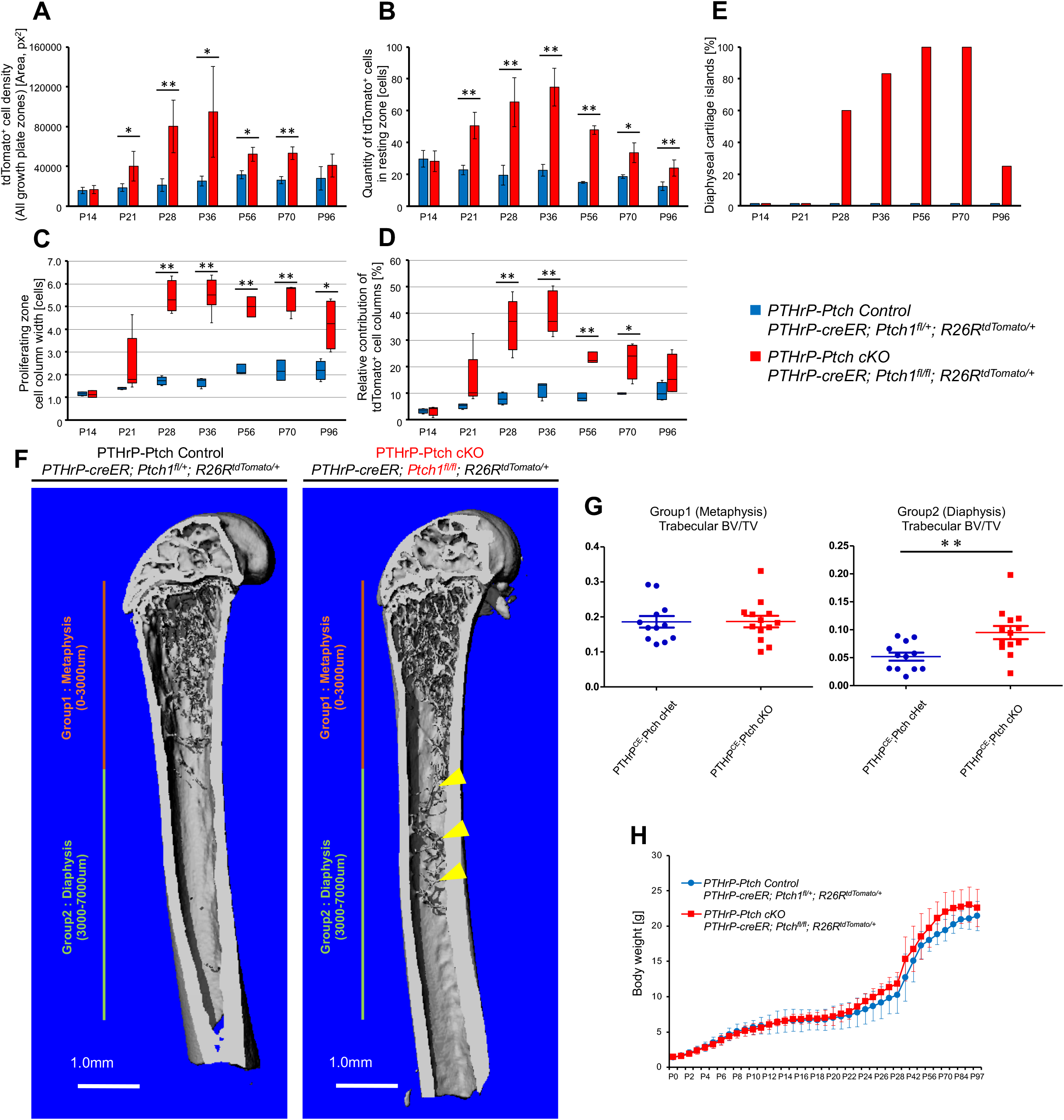
Transient clonal competency of Hedgehog-activated PTHrP^+^ resting chondrocytes and their contribution to trabecular bone formation. **(A-E)** Cell quantification. (A): tdTomato^+^ lineage-marked cells in growth plate. Image J “Measure” function was utilized to quantify tdTomato^+^ cells across all zones of the growth plate as a function of area (px^2^). (B): tdTomato^+^ lineage-marked cells in resting zone. Images were processed in Image J to create a selective mask excluding all cells outside of this zone prior to quantification. (C): Lateral cell width of columns in the proliferating zone manually counted at each time point. (D): Relative contribution of tdTomato^+^ columns across growth plate. A grid overlay function in Image J software was used to assist assessment of cell columns across the width of the growth plate. Number of tdTomato^+^ chondrocyte columns was summed and calculated as a percentage of the total number of cell columns (including tdTomato^neg^ columns) across the lateral width of the growth plate. (E): Graph representation of the percentage of femurs containing at least one histological section with visible cartilage islands in diaphysis. Blue bars: PTHrP-Ptch Control cells, red bars: PTHrP-ΔPtch cells. P14: *n*=4 (Control), *n*=4 (PTHrP-Ptch cKO). P21: *n*=4 (Control), n=5 (PTHrP-Ptch cKO). P28: *n*=4 (Control), *n*=5 (PTHrP-Ptch cKO). P36: *n*=4 (Control), *n*=6 (PTHrP-Ptch cKO). P56: *n*=3 (Control), n=3 (PTHrP-Ptch cKO). P70: *n*=3 (Control), *n*=4 (PTHrP-Ptch cKO). P96: *n*=4 (Control), *n*=4 (PTHrP-Ptch cKO). \**p*<0.05, \*\**p*<0.01, two-tailed, Mann-Whitney’s *U*-test. Data are presented as mean ± s.d. **(F,G)** 3D-μCT analysis. (F): Representative 3D-μCT images of *PTHrP-creER; Ptch1^fl/+^; R26R^tdTomato^* (PTHrP-Ptch Control, A) and *PTHrP-creER; Ptch1^fl/fl^; R26R^tdTomato^* (PTHrP-Ptch cKO, B,C) femurs at P96 (pulsed at P6). Arrowheads: ectopic trabecular bone, Group 1: metaphysis (0–3000 μm) and Group 2: diaphysis (3000–7000 μm). Scale bars: 1 mm. (G): Trabecular BV/TV in Group 1 (metaphysis) (left) and Group 2 (diaphysis) (right). *n*=12 (Control), *n*=13 (PTHrP-Ptch cKO) mice, including both sexes. Scale bars: 100 μm. \*\**p*<0.01, two-tailed, Mann-Whitney’s *U*-test. Data are presented as mean ± s.d. **(H)** Body weight, from P0 to P97. Blue line: PTHrP-Ptch Control, red: PTHrP-Ptch cKO mice.

We further examined the cell width of the tdTomato^+^ columns within the proliferating zone. PTHrP^CE^-P6 Control cells established columns with ∼1 cell in width at P14 and P21, then columns with ∼2 cells in width from P28 to P96 (Fig.4C, blue bars). In contrast, in the PTHrP-Ptch cKO, PTHrP^CE^ΔPtch-P6 cells established significantly wider columns starting from P28: the cell width of PTHrP^CE^ΔPtch-P6 columns increased drastically between P21 and P28, reaching a maximum average of 5-6 cells in width at P36 (Fig.4C, red bars), though this was associated with significant variability at all time points after P14. These findings support the notion that Hedgehog-activated PTHrP^CE^ΔPtch-P6 resting chondrocytes can establish larger columns of chondrocytes, contributing to the observed clonal competency and hyperplasia.

We further enumerated the relative contribution of tdTomato^+^ cell columns among all columns within the growth plate. In the PTHrP-Ptch Control growth plate, PTHrP^CE^-P6 columns contributed to approximately ∼10% of the entire columns at and after P28 (Fig.4D, blue bars). In contrast in the PTHrP-Ptch cKO growth plate, PTHrP^CE^ΔPtch-P6 columns increased substantially between P14 and P36, reaching a maximum contribution of ∼40% at P36 which decreased toward P96 (Fig.4D, red bars). These quantitative results corroborate the histological observation that PTHrP^CE^ΔPtch-P6 resting chondrocytes transiently acquire clonal dominance which is normalized in later stages.

Our histological observation in Figure 3 demonstrates that some of the PTHrP^CE^ΔPtch-P6 chondrocytes can separate from the hyperplastic growth plate and migrate toward the marrow space, ectopically forming discrete cartilage islands. We therefore quantified the occurrence of these ectopic cartilage islands within the marrow space. Importantly, no cartilage islands were observed in PTHrP-Ptch Control mice at any time point (Fig.4E, blue bars). In contrast, in the PTHrP-Ptch cKO, cartilage islands first appeared at P28 and increased in prevalence over time, with 100% of all PTHrP-Ptch cKO samples possessing cartilage islands at P56 and P70 (Fig.4E. red bars). However, interestingly, these tdTomato^+^ cartilage islands rapidly disappeared by P96, indicating that these islands are not permanent. Of note, these cartilage islands are generally located in proximity to the growth plate and later migrate toward the diaphysis over time.

The disappearance of the ectopic cartilage islands at P96 prompted us to examine quantitively if these cartilaginous structures can transform into mineralized bone structures. To this end, we performed a 3D microCT analysis of PTHrP-Ptch Control and PTHrP-Ptch cKO bones at P96 (Fig.4F). We measured the trabecular bone volume per tissue volume (BV/TV) of two specific locations: Group 1 (Metaphysis) in the area from 0 to 3 mm away from the growth plate, and Group 2 (Diaphysis) in the area from 3 to 7 mm away from the growth plate. Importantly, no statistically significant difference in the trabecular BV/TV was observed between PTHrP-Ptch Control and PTHrP-Ptch cKO bones in the metaphysis (Group 1) (Fig.4G, left panel). However, the trabecular BV/TV of the diaphysis (Group 2) was significantly increased in the PTHrP-Ptch cKO (Fig.4G, right panel), demonstrating that chondrocyte-to-osteoblast differentiation occurred in this area and resulted in mineralized tissue formation (see also Suppl. Fig.2A,B). Indeed, apparently contiguous structures of bone trabeculae were observed in the PTHrP-Ptch cKO diaphyseal marrow (Fig.4F, yellow arrowheads), which may represent traces of cartilages island transforming into mineralized structures. No difference was observed in the cortical BV/TV between the two groups (Suppl. Fig.2C). No significant difference was observed in body weight between the two groups at any time point (Fig.4H). Furthermore, the bone length of the PTHrP-Ptch cKO femur was equivalent to that of the PTHrP-Ptch Control femur at all time points except in earlier time points of P14 and P21 (Suppl. Fig.2D), indicating that Hedgehog activation negatively affects the bone length only transiently. Therefore, the specific increase in trabecular bone mass is unlikely to be attributable to changes in systemic factors or overall bone morphology.

## Discussion

Here in this study, we identified that PTHrP^+^ chondrocytes in the resting zone of the growth plate can give rise to trabecular bone-forming osteoblasts within the marrow space, when Hedgehog signaling is specifically activated in these cells (see concluding diagram, Fig.5). The process of resting chondrocyte-to-trabecular bone osteoblast transformation is mediated by transient amplification of proliferating chondrocytes, which might represent transit-amplifying precursors for trabecular bone-forming osteoblasts. The findings from our in vivo functional genetic cell lineage analysis provide definitive supporting evidence for two important processes regulating bone formation. First, hypertrophic chondrocytes can transform into osteoblasts and contribute to trabecular bone formation, and second, Hedgehog activation induces a common outcome of promoting osteoblast differentiation of early precursor cells, i.e. growth plate chondrocytes. Paracrine signals by Indian hedgehog (Ihh) released from the prehypertrophic zone may exert effects not only on cells in the adjacent perichondrium, but also on chondrocytes in the adjacent proliferating zone to facilitate their osteogenic differentiation, therefore engaging multiple cellular sources to achieve endochondral bone formation (35).

**Figure 5.**
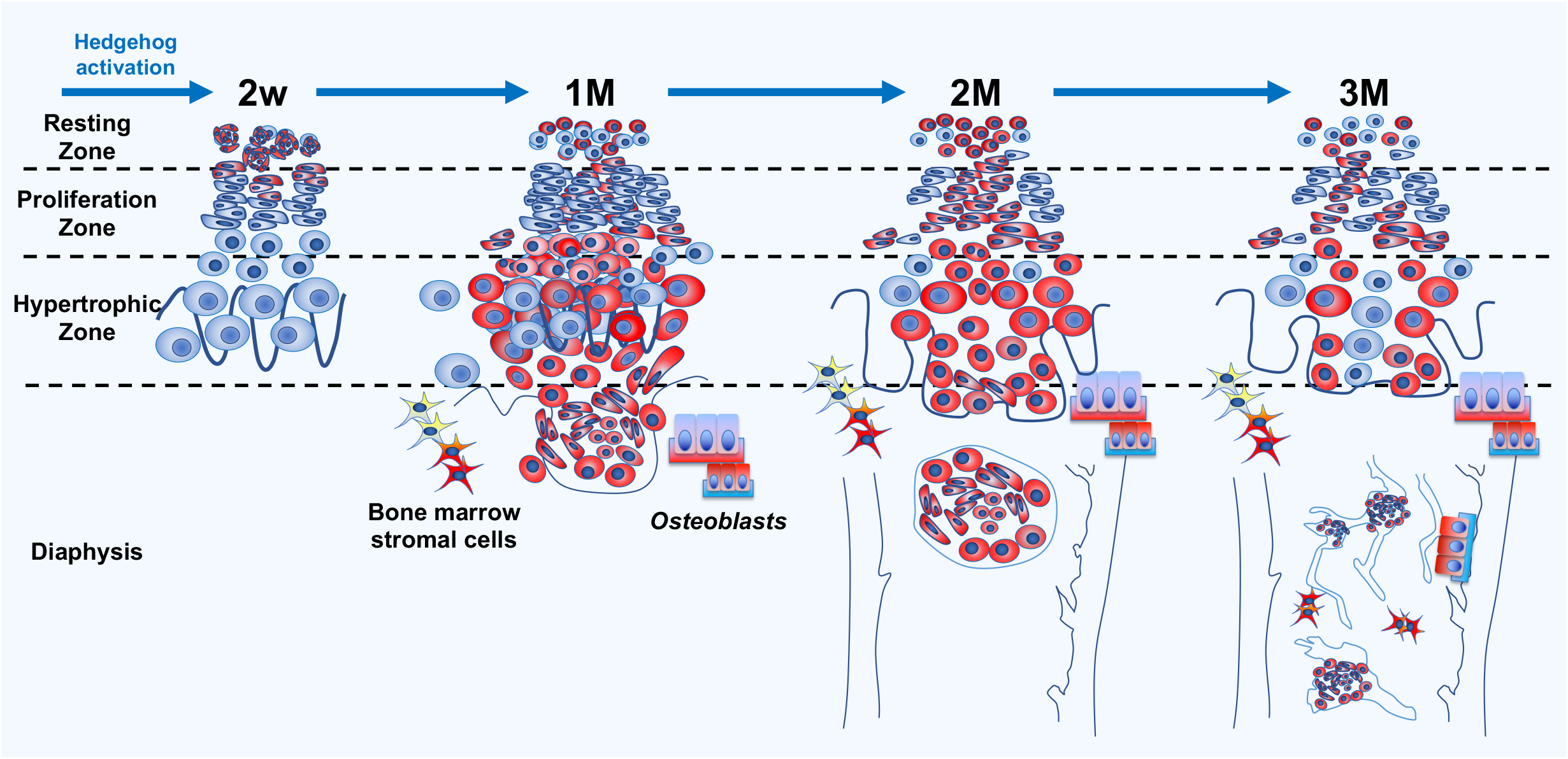
Growth plate resting zone chondrocytes acquire transient clonal competency upon hedgehog activation and transform into trabecular bone osteoblasts. In *PTHrP-creER; Ptch1^fl/fl^; R26R^tdTomato/+^* (PTHrP-Ptch cKO) mice, only a subset of cells in RZ become activated for Hedgehog signaling upon tamoxifen injection. Cells marked by tdTomato are surrounded by tdTomato-negative cells. Therefore, our resting chondrocyte-specific *PTHrP-creER* line allows a low mosaicism analysis. PTHrP^CE^ΔPtch-P6 tdTomato^+^ cells in PTHrP Ptch-cKO mice clonally expand and form “*patched roses*” after two weeks. Significant growth plate hyperplasia and distortion are visible after one month after pulse, and then PTHrP^CE^ΔPtch-P6 tdTomato^+^ cells separate from the growth plates and migrate to the bone marrow after two months. Growth plate hyperplasia in PTHrP-Ptch cKO mice decreases to about the same level as in Control mice. In addition, many PTHrP^CE^ΔPtch-P6 tdTomato^+^ cells present in bone marrow and on trabecular surfaces in PTHrP-Ptch cKO mice associated with an increase of trabecular bones.

We observed that Hedgehog activation transiently promotes clonal competency of PTHrP^+^ resting chondrocytes within the growth plate, resulting in the dominance of Hedgehog-activated clones during a defined period. Interestingly, Hedgehog-activated chondrocytes in the resting zone undergo an abrupt increase in clonality between two to three weeks postnatally, followed by a gradual decrease in clonal competency thereafter, indicating that a cell signaling “*switch*” might occur during these periods. While it is likely that excessive proliferation of Hedgehog-activated chondrocytes is at least in part a function of cell-autonomous effects, we assume that an external physiological cue plays a role in the exaggerated proliferative response of these cells in a defined period. In the absence of such exogenous signals, we might expect Hedgehog-activated resting chondrocytes to be present in greater quantities at P14, a finding which we did not observe. Signaling switches occurring during growth plate development might exert some levels of control over the response of chondrocytes in the presence of Hedgehog activation, presumably through activation of parallel and synergistic pathways or through regulation of shared downstream inhibitors. While Hedgehog-activated chondrocytes are prone to excessive proliferation at earlier time points, these cells show neither resistance to apoptosis or inhibition of hypertrophy. The response of growth plate chondrocytes to Hedgehog activation appears to be context-dependent without being restricted to a canonical action to delay chondrocyte hypertrophy or inhibit differentiation (36, 37). Our results also support a hypothesis that Hedgehog activation in PTHrP^+^ resting chondrocytes induces only a transient increase in proliferative activities, presumably due to restraint on signaling by external cues. Dissecting detailed molecular interactions between Hedgehog and other major signaling pathways in regulating skeletal stem cell behaviors within the resting zone presents an important opportunity for future studies.

Interestingly, Hedgehog activation in only a small number of PTHrP^+^ chondrocytes within the resting zone is sufficient to induce a cell-autonomous increase in the trabecular bone in a defined area of the diaphyseal marrow space. It is reasonable to presume chondrocyte-to-osteoblast differentiation stems from direct downstream effects of Hedgehog signaling, indirect feedback from interdependent pathways, or most likely, a combination of both. Our study demonstrates for the first time the link between PTHrP^+^ skeletal stem cells in the resting zone and trabecular bone osteoblasts in the diaphyseal marrow space. It will be interesting to examine in future studies how Hedgehog signaling spatiotemporally regulates a small number of stem cells within the resting zone and eventually exerts impact on trabecular bone formation under physiological and pathological conditions.

In conclusion, our findings collectively demonstrate that Hedgehog activation drives resting zone chondrocytes into transit-amplifying states as proliferating chondrocytes and eventually converts these cells into osteoblasts, unraveling a novel Hedgehog-mediated mechanism that facilitates osteogenic cell fates of PTHrP^+^ skeletal stem cells. The findings from our in vivo genetic functional cell lineage analysis provide a solid foundation to understand how therapeutic intervention to the Hedgehog signaling pathway may impact a group of stem cell populations residing in the resting zone of the growth plate of growing individuals and subsequently affect bone formation.

## Methods

### Mice

*PTHrP-creER* mice have been previously (6). *Rosa26-CAG-loxP-stop-loxP-*tdTomato (Ai14: *R26R*-tdTomato, JAX007914) mice, *Ptch1*-floxed mice (JAX012457) and *Smo*-floxed (JAX004526) mice were acquired from the Jackson laboratory. All procedures were conducted under the University of Texas Health Science Center at Houston’s Animal Welfare Committee (AWC), protocol AWC-21-0070, and the University of Michigan’s Institutional Animal Care and Use Committee (IACUC), protocols 5703 and 7681. All mice were housed in a specific pathogen-free condition and analyzed in a mixed background. Mice were housed in static microisolator cages (Allentown Caging, Allentown, NJ). Access to water and food (irradiated LabDiet 5008, Richmond, IN) was ad libitum. Animal rooms were climate controlled to provide temperatures of 22-23°C on a 12 h light/dark cycle (lights on at 0600). For all breeding experiments, *creER* transgenes were maintained in male breeders to avoid spontaneous germline recombination. Mice were identified by micro-tattooing or ear tags. Tail biopsies of mice were lysed by a HotShot protocol (incubating the tail sample at 95°C for 30 min in an alkaline lysis reagent followed by neutralization) and used for PCR-based genotyping (GoTaq Green Master Mix, Promega, and Nexus X2, Eppendorf). Perinatal mice were also genotyped fluorescently (BLS miner’s lamp) whenever possible. Mice were euthanized by over-dosage of carbon dioxide or decapitation under inhalation anesthesia in a drop jar (Fluriso, Isoflurane USP, VetOne).

### Tamoxifen and induction of cre-loxP recombination for PTHrP-creER line

Tamoxifen (Sigma T5648) was mixed with 100% ethanol until completely dissolved. Subsequently, a proper volume of sunflower seed oil (Sigma S5007) was added to the tamoxifen-ethanol mixture and rigorously mixed. The tamoxifen-ethanol-oil mixture was incubated at 60°C in a chemical hood until the ethanol evaporated completely. The tamoxifen-oil mixture was stored at room temperature until use. Treatment of *PTHrP-creER; Ptch1^fl/+^; R26R^tdTomato^*and *PTHrP-creER; Ptch1^fl/fl^; R26R^tdTomato^* mice with various doses of tamoxifen at P6 and P9 was performed previously and determined to produce optimal responsiveness of the reporter allele at P6. Therefore, we consistently induced the lineage at P6 throughout this study.

For the EdU assay*, PTHrP-creER; Ptch1^fl/+^; R26R^tdTomato^* and *PTHrP-creER; Ptch1^fl/fl^; R26R^tdTomato^* mice received 500µg of tamoxifen intraperitoneally at P6, followed by two serial injections of EdU (200µg) at a 3-hour interval shortly before analysis at P21.

### Histology and Immunohistochemistry

Samples were dissected under a stereomicroscope (Nikon SMZ-800) to remove soft tissues, fixed in 4% paraformaldehyde at 4°C, then decalcified in 15% EDTA for up to 14 days. Decalcified samples were cryoprotected in 30% sucrose/PBS solutions and then in 30% sucrose/PBS:OCT (1:1) solutions, each at least overnight at 4°C. Samples were embedded in an OCT compound (Tissue-Tek, Sakura) under a stereomicroscope and transferred on a sheet of dry ice to solidify the compound. Embedded samples were cryosectioned at 50μm using a cryostat (Leica CM1850) and adhered to positively charged glass slides (Fisherbrand ColorFrost Plus). Cryosections were stored at −20°C until use. Sections were postfixed in 4% paraformaldehyde for 15 min at room temperature. For immunostaining, sections were permeabilized with 0.25% TritonX/TBS for 30 min, blocked with 3% bovine serum albumin/TBST for 30 min. and incubated with goat anti-osteopontin (OPN) polyclonal antibody (1:500, R&D, AF808), rabbit anti-collagen 1 (Col1) polyclonal antibody (1:500, Cedarlane, CL50151AP) and rabbit anti-Caspase-3 polyclonal antibody (1:250, Promega/Fisher, PR-G7481) overnight at 4 °C, and subsequently with Alexa Fluor 647-conjugated donkey anti-goat IgG (A21082), or Alexa Fluor 647-conjugated donkey anti-rabbit IgG (A31573) (1:400, Invitrogen) for 3 h at room temperature. For EdU assays, sections were incubated with Alexa Fluor 488-azide (Invitrogen A10266) for 30 min at 43°C using Click-iT Imaging Kit (Invitrogen, C10337). Sections were further incubated with DAPI (4’,6-diamidino-2-phenylindole, 5µg/ml, Invitrogen D1306) to stain nuclei prior to imaging. Stained samples were mounted in TBS with No.1.5 coverslips (Fisher).

### Imaging

Images for fixed sections were captured by an automated inverted fluorescence microscope with a structured illumination system (Zeiss Axio Observer Z1 with ApoTome.2 system) and Zen 2 (blue edition) software. Images were typically tile-scanned with a motorized stage, Z-stacked and reconstructed by a maximum intensity projection (MIP) function. The filter settings used were: FL Filter Set 31 (Ex. 565/30, Em. 620/60 nm), Set 34 (Ex. 390/22, Em. 460/50 nm), Set 38 HE (Ex. 470/40, Em. 525/50 nm), Set 43 HE (Ex. 550/25, Em. 605/70 nm), Set 46 HE (Ex. 500/25, Em. 535/30 nm), Set 47 HE (Ex. 436/20, Em. 480/20 nm), Set 50 (Ex. 640/30, Em. 690/50 nm) and Set 63 HE (Ex. 572/25, Em. 629/62 nm). The objectives used were: Fluar 2.5x/0.12, EC Plan-Neofluar 5x/0.16, Plan-Apochromat 10x/0.45, EC Plan-Neofluar 20x/0.50, EC Plan-Neofluar 40x/0.75, Plan-Apochromat 63x/1.40. Differential interference contrast (DIC) was used for objectives higher than 10x. Representative images of at least three independent biological samples are shown in all figures.

### Image Quantification

NIH ImageJ (Fiji, v1.52i) software was used for image manipulation and quantification. For cell quantification, serial sections (15∼20 sections of 25∼50µm thickness each, total 375∼1,000µm thickness) were evaluated for each independent sample. A minimum of *n*=4 femurs for both Control and PTHrP-Ptch cKO groups at each time point were analyzed, and values for each of appx. 20 sections per femur were averaged where appropriate.

Images of histological sections obtained using Zen 2 (blue edition) image capture software yielded raw TIFF files with 96 dpi horizontal and vertical resolution and a bit depth of 24. Images were imported into ImageJ and a JAVA macro was developed to allow batch processing; image thresholding was performed to select for red (tdTomato^+^) cells, and color was subsequently set to binary. All images were cropped to an identical size of 3,960,000 pixels^2^ using a manually set rectangular ROI centered around the growth plate. ImageJ “Watershed” function was utilized to improve resolution by dividing large cell clusters into discrete particles.

A. For quantification of cell density among all growth plates zones, the ImageJ “Measure” function was utilized to determine the value of black pixels on a white background subsequent to color thresholding. Results were measured in px^2^.
B. For quantification of cell density limited to the resting zone, images were further processed to create a selective ROI excluding all cells outside of this zone. A freehand ROI was manually set for each section and a mask was created to limit quantification to resting zone cells.
C. For analysis of proliferating zone cell column width, manual count of the maximum number of cells in the widest cell column (perpendicular to longitudinal axis of femur) per section was performed and results were subsequently averaged.
D. For analysis of relative percentage of labeled cell columns across the growth plate, four sections were selected per sample (sections 5, 10, 11, and 15 in numerical order) and imported to ImageJ. A grid overlay with horizontal and vertical lines spaced 20 pixels apart was applied to each image, such that one square cell of the grid approximated the size of one cell within the proliferating zone. Number of tdTomato^+^ columns within the grid was summed and values representing the proportion of tdTomato^+^ columns out of total number of cell columns (including tdTomato^-^ columns) across the width of the growth plate was displayed as a percentage.
E. For analysis of migratory *patched roses*, all sections for both Ptch cHet and cKO samples at each time point were evaluated for either the presence or absence of lesions. If *patched roses* were observed either within the growth plate or marrow space of at least one section of a femur, that sample was deemed positive for the presence of migratory lesions.

### Statistical Analysis

Non-parametric evaluation between experimental and control samples at each timepoint and for each separate parameter of data quantification (A-D above) was performed using a Mann-Whitney’s *U* test. A *p*-value of <0.05 was considered significant (*Table I*). Intra-examiner reliability (repeatability) for each method of data assessment was evaluated by repeating quantification on one half of samples selected at random and comparing initial and repeated outcomes using Intraclass Correlation Coefficient (ICC) calculations (*Table IV*).

Statistical advisement was obtained from UM Consulting for Statistics, Computing and Analytics Research (CSCAR). Sample size was determined on the basis of previous literature and our previous experience to give sufficient standard deviations of the mean. Investigators were not blinded during experiments and outcome assessment. Mice were used regardless of the sex. All data and images are representative of a minimum of three independent biological samples.

## Acknowledgements

We thank S.Y. Wong for *Ptch1*-floxed mice. This research was supported by National Institute of Health grants R01DE026666, R01DE030630 (to N.O.), R01DE029181 (to W.O.), and University of Michigan MCubed 2.0 Grant (to N.O., W.O.).

---

**Figure S1.**
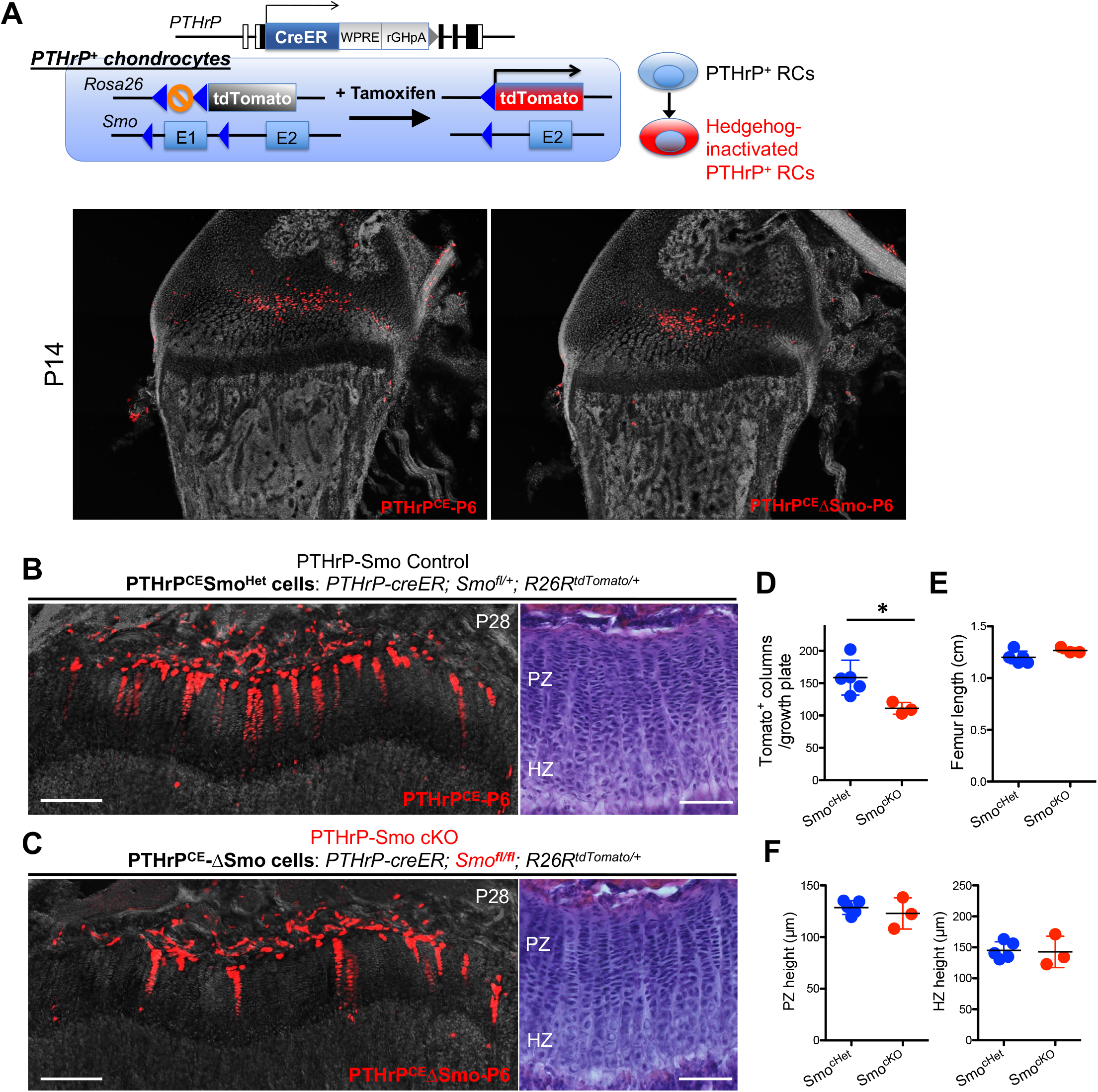
Effects of Hedgehog inactivation on columnar chondrocyte formation of PTHrP^+^ resting chondrocytes. **(A)** Experimental design to analyze cell fates of Hedgehog-inactivated PTHrP^+^ resting chondrocytes. Triple transgenic mice carrying *PTHrP-creER*, *R26R-*tdTomato allele and *Smoothened (Smo)*-floxed allele received a single dose of tamoxifen at P6. In PTHrP^+^ chondrocytes, the stop cassette of the *Rosa26* locus and exon 1 of the *Smo* locus are simultaneously removed upon tamoxifen injection. As a result of this recombination, Hedgehog-inactivated cells can be traced by tdTomato in *PTHrP-creER*; *Smo*^fl/fl^; *R26R*^tdTomato/+^ mice. **(B, C)** *PTHrP-creER; Smo^fl/+^; R26R^tdTomato/+^* (PTHrP-Smo Control, B) and *PTHrP-creER; Smo^fl/fl^; R26R^tdTomato/+^* (PTHrP-Smo cKO, C) distal femur growth plates at P28, pulsed at P6. Red in (B): PTHrP^CE^Smo^Het^-tdTomato^+^ cells. Red in (C): PTHrP^CE^ΔSmo-tdTomato^+^ cells. Right panels: H&E staining. PZ: proliferating zone, HZ: hypertrophic zone. Scale bars: 200µm (left), 100µm (right). **(D-F)** Quantification. The number of tdTomato^+^ columns in growth plates (D), femur length (E) and the height of proliferating (F, left panel) and hypertrophic zone (F, right panel). *n*=5 mice for PTHrP-Smo Control, *n*=3 mice for PTHrP-Smo cKO. \**p*<0.05, Mann-Whitney’s *U*-test.

**Figure S2.**
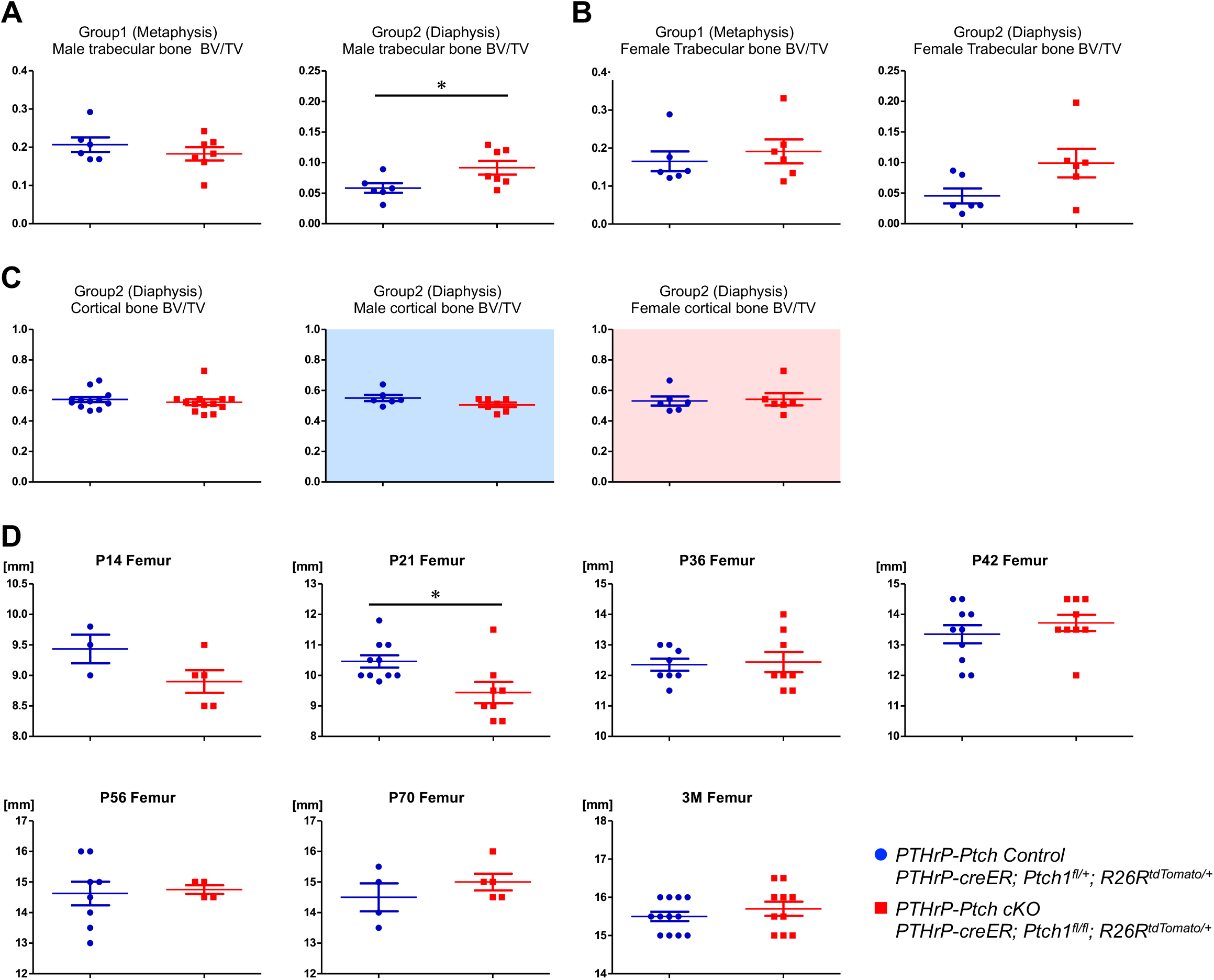
Three-dimensional microCT analysis of PTHrP^+^ Ptch1 conditional mutant bone. **(A-C)** 3D-μCT analysis of *PTHrP-creER; Ptch1^fl/+^; R26R^tdTomato^* (PTHrP-Ptch Control, A) and *PTHrP-creER; Ptch1^fl/fl^; R26R^tdTomato^* (PTHrP-Ptch cKO, B,C) femurs at P96 (pulsed at P6). Blue: Control, red: PTHrP-Ptch cKO. (A,B): Trabecular bone BV/TV in male (A, *n*=6) and female (B, *n*=6) mice, in Group 1 (metaphysis) (left) and Group 2 (diaphysis) (right). (C): Cortical bone BV/TV in Group 2 (diaphysis), including both sexes. *n*=12 (Control), *n*=13 (PTHrP-Ptch cKO) mice, including both sexes. Scale bars: 100 μm. \**p*<0.05, two-tailed, Mann-Whitney’s *U*-test. Data are presented as mean ± s.d. **(D)** Femur length of *PTHrP-creER; Ptch1^fl/+^; R26R^tdTomato^* (PTHrP-Ptch Control, A) and *PTHrP-creER; Ptch1^fl/fl^; R26R^tdTomato^* (PTHrP-Ptch cKO, B,C) femurs at P96 (pulsed at P6), at P14, 21, 36, 42, 56, 70, 96. Blue: Control, red: PTHrP-Ptch cKO. \**p*<0.05, two-tailed, Mann-Whitney’s *U*-test. Data are presented as mean ± s.d.

